# CD20 expression regulates CD37 levels in B-cell lymphoma – implications for immunotherapies

**DOI:** 10.1101/2023.12.06.570441

**Authors:** Malgorzata Bobrowicz, Aleksandra Kusowska, Marta Krawczyk, Aleksander Slusarczyk, Joanna Barankiewicz, Joanna Domagala, Matylda Kubacz, Michal Šmída, Lenka Dostalova, Katsiaryna Marhelava, Klaudyna Fidyt, Christopher Forcados, Monika Pepek, Iwona Baranowska, Anna Szumera-Cieckiewicz, Else Marit Inderberg, Sébastien Wälchli, Agnieszka Graczyk-Jarzynka, Carina Lynn Gehlert, Matthias Peipp, Malgorzata Firczuk, Monika Prochorec-Sobieszek, Magdalena Winiarska

## Abstract

Rituximab (RTX) plus chemotherapy (R-CHOP) applied as a first-line therapy for lymphoma leads to a relapse in approximately 40% of patients. Therefore, novel approaches to treat aggressive lymphomas are being intensively investigated. Several RTX-resistant (RR) cell lines have been established as surrogate models to study resistance to R-CHOP. Our study reveals that RR cells are characterized with a major downregulation of CD37, a molecule currently explored as a target for immunotherapy. Using CD20 knockout (KO) cell lines, we demonstrate for the first time that CD20 and CD37 form a complex and the presence of CD20 stabilizes CD37 in the cell membrane. Consequently, we observe a diminished cytotoxicity of anti-CD37 monoclonal antibody (mAb) in complement-dependent cytotoxicity in both RR and CD20 KO cells that can be partially restored upon lysosome inhibition. On the other hand, the internalization rate of anti-CD37 mAb in CD20 KO cells is increased when compared to controls, suggesting unhampered efficacy of antibody drug conjugates. Importantly, even a major downregulation in CD37 levels does not hamper the efficacy of CD37-directed chimeric antigen receptor (CAR) T cells. In summary, we present here a novel mechanism of CD37 regulation with further implications for the use of anti-CD37 immunotherapies.

## Introduction

Diffuse large B-cell lymphoma (DLBCL) is the most frequent non-Hodgkin lymphoma (NHL) subtype, accounting for about 40% of NHL cases and is also one of the most aggressive subtypes(1). For years, the first-line therapy in DLBCL has been R-CHOP – a combination of chemotherapeutics with rituximab – an anti-CD20 monoclonal antibody (mAb). Although the treatment is mostly well-tolerated and efficient, approximately 40% of patients face relapse characterized by a very poor prognosis and the median survival following relapse not exceeding 6 months(2). In the recent few years, considerable progress has been made in the registration of novel therapies used as a second-line treatment, that nowadays force the immunotherapeutic approaches like antibodies and antibody drug-conjugates directed against CD19 or CD79b molecules as well as CD19 CAR-T cell formulations - especially dedicated for primary refractory patients. Numerous clinical trials are currently underway exploring new molecular targets against aggressive lymphomas including those testing CD37-directed immunotherapies (NCT04136275, NCT04358458).

The molecular changes induced by RTX and leading to a failure of the next-line therapies have been widely studied in established B-cell lymphoma cell lines with induced resistance to RTX (3–5). These models confirmed CD20 downregulation already reported in R-CHOP – treated patients (6–8). In this study, we have sought to thoroughly characterize the RTX-resistant cell lines to better understand their phenotype. Our analyses revealed a significant downregulation of CD37 molecule on the surface of RR cells. CD37, a tetraspanin protein with structural similarity to CD20, is expressed exclusively on hematopoietic cells and its highest levels are detected in the B-cell lineage from pre-B to mature B cells (9,10). Alike CD20, CD37 is absent on the surface of plasma cells, making it a suitable target for immunotherapy. In fact, numerous anti-CD37 targeting immunotherapies are currently under investigation (11–14). Besides being a molecular target for immunotherapies, CD37 is an important player in the biology of B-cell malignancies (15) as demonstrated in murine models, where CD37 has been described as a negative regulator of lymphomagenesis (16). Recently, CD37 has been reported to inhibit the fatty acids (FA) transporter FATP1, so CD37-negative lymphoma cells take advantage of increased FA uptake and process exogenous palmitate into energy supporting their increased proliferation (17). In line with that, CD37 presence on the cell surface has been shown to correlate with overall survival (OS) and progression-free survival (PFS) in DLBCL patients (18). As patients with high CD37 expression showed improved survival on R-CHOP regardless of *CD20* expression (18), CD37 has been suggested to act as a “molecular facilitator” of rituximab action in a yet undefined mechanism. Intriguingly, surface CD37 levels have been shown to correlate with CD20 levels as assessed by flow cytometry in DLBCL cell lines(18). In this study, using RR and CD20 KO cell lines, we investigated the role of CD20 in regulating CD37 levels and its consequences for the efficacy of mAbs and CAR-based immunotherapy.

## Materials and methods

### Cell culture

All cell lines were cultured at 37°C in a fully humidified atmosphere of 5% CO2 in RPMI-1640 (Invitrogen), supplemented with 10% heat-inactivated fetal bovine serum (FBS), 100U/mL of penicillin and 100 μg/mL of streptomycin. Cells were passaged every other day. Rituximab-resistant cell lines and their wild-type counterparts were kindly provided by Prof. F.Hernandez-Ilizaliturri from Roswell Park, NY, US (Raji, RL, U2932 cell lines) and by Dr. Michal Šmída from the Central European Institute of Technology, Czech Republic (Ramos cell lines). OCI-Ly-7 (named LY7 in this paper), SU-DHL4 (further referred to as DHL4) cell lines and HEK-293T used for lentivirus production were purchased from DSMZ. Daudi cells were purchased from ATCC.

### Retroviral T cells transduction

Peripheral blood mononuclear cells (PBMCs) were isolated from buffy coats from healthy donors by density gradient centrifugation using Lymphoprep™ (STEMCELL Technologies Canada, Inc.). This procedure was approved by the Bioethics Committee of the Medical University of Warsaw, Poland. Following isolation, PBMCs were seeded onto 6-well plate at density 2 × 10^6^ cells/mL in full RPMI-1640 medium and stimulated for 48h with anti-CD3 (1:1000) and anti-CD28 (1:1000) mAbs (Invitrogen). After 48h, stimulated PBMCs were collected and seeded onto a 24-well plate coated with 50 μg/ml retronectin (TakaraBio) at a density 1 × 10^6^ cells/mL. For retroviral transduction, two rounds of spinoculation with non-concentrated retroviral supernatants were performed (1h, 1250 rpm, 32°C). Six hours following the second spinoculation, viral supernatants were replaced with full RPMI-1640 medium supplemented with 200 U/ml IL-2 (PeproTech) and Dynabeads Human T-Activator CD3/CD28 (Thermo Fisher Scientific).

### PLA assay

Duolink^®^ Proximity Ligation Assay kit (Sigma Aldrich) was used to determine the interaction between CD20 and CD37. Briefly, Raji and U2932 NTC and CD20 KO cells were incubated with anti-CD37 mouse Ab (M-B371 clone, BD) and anti-CD20 human Ab (obinutuzumab, Roche). Relevant negative controls were performed using isotype antibodies and cells incubated with anti-CD20 human and anti-CD20 mouse antibody served as a positive control. Afterwards, the cells were incubated with anti-human and anti-mouse probes followed by in situ ligation and amplification according to the manufacturer’s protocol. Duolink^®^ Green Detection kit was used to detect protein interactions and the cells were analysed on BD Fortessa.

### Immunohistochemistry of primary DLBCL samples

Two tissue microarrays were prepared from archival formalin-fixed, paraffin-embedded (FFPE) blocks from DLBCL patients diagnosed in the Institute of Hematology and Transfusion Medicine. From each case, two 1 mm cores were evaluated. The slides were stained with anti-CD37 (clone 2B8; Thermo Fisher Scientific, pH 6.0, 1:200), anti-CD20 and anti-CD19 mAbs (Dako, RTU). The positive controls included lymphoid tissue in the tonsil, appendix, and spleen. CD37 surface expression was scored as negative (<5%) and positive (≥5%) as previously reported (18), CD20 and CD19 surface were positive according to criteria of WHO Classification of Tumours of Haematopoietic and Lymphoid Tissues 2017.

### Patients’ sample analysis

We retrospectively analyzed 47 patients diagnosed with DLBCL in the Institute of Hematology and Transfusion Medicine, Warsaw, Poland. Following patients’ diagnosis data were obtained: age, sex, disease stage according to Ann Arbor classification, treatment response determined by Lugano criteria (19), progression-free and overall survival (PFS and OS, respectively). All patients received R-CHOP regimen as first-line treatment. The data collection was conducted according to the Declaration of Helsinki, and the protocol was approved by the Ethics Committee of the Institute of Hematology and Transfusion Medicine, Warsaw, Poland.

### Statistical analysis

Results were plotted with GraphPad Prism. Statistical significance was assessed by appropriate tests provided in figure legends. The *p*-values were marked with asterisks on the charts (\**p* <0.05, \*\**p* <0.01, \*\*\**p* <0.001, \*\*\*\**p* <0.0001).

## Results

### Decrease in CD20 is a hallmark of RR cells

RR cells provide an in vitro model for investigating the consequences of prolonged exposition of tumor cells to rituximab (4,20). As already reported by others (4,20), CD20 downregulation is a hallmark of RR cells (Fig. 1A), which largely contributes to their resistance to RTX in complement-dependent cytotoxicity (CDC) (Fig. 1B). Moreover, these cells are resistant to RTX in vivo as assessed by mice survival (Suppl. Fig. 1A) and tumor burden (Suppl. Fig. 1B). To investigate the biological consequences of CD20 decrease, we generated models of CD20 KO in several B-cell lymphoma cell lines using CRISPR-Cas9 technology (Fig. 1C). All generated CD20 KO cells were characterized with resistance to rituximab (Fig. 1D).

**Figure 1.**
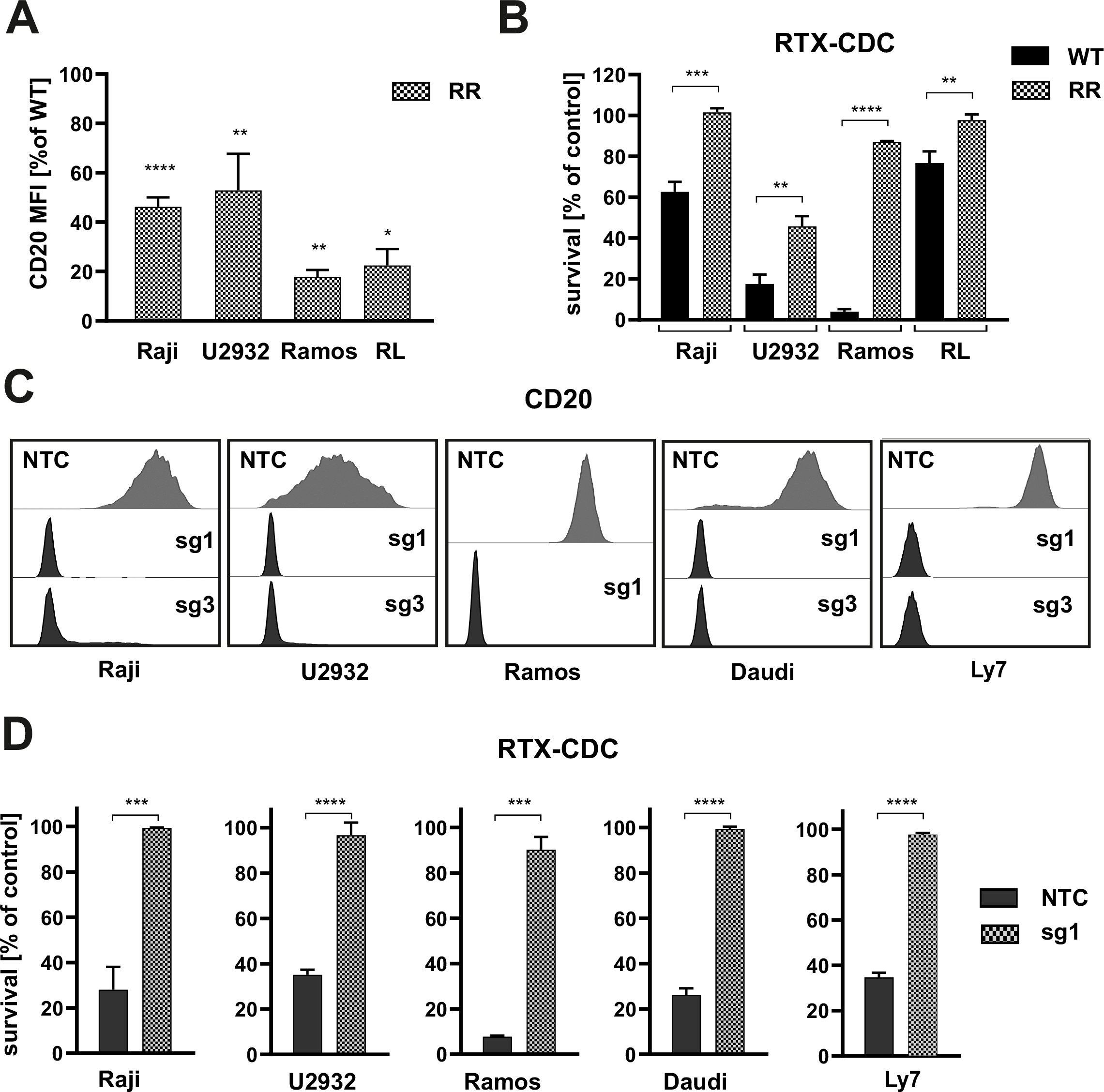
Decrease in CD20 is a hallmark of RTX-resistant cells. **(A)** RTX-resistant (RR) cells were stained with FITC-conjugated anti-CD20 mAb. Dead cells were dsicriminated upon staining with PI. The results are presented as a percentage of mean fluorescence intensity (MFI) of WT cells (mean± SEM). Statistical significance was determined with Welch’s *t*-test, \**p<*0.05, \*\**p*<0.01, \*\*\*\**p*<0.0001 vs controls. **(B)** Equal amounts of WT and RR cells were incubated for 1 h (Raji, Ramos) or 4h (U2932, RL) with 100 μg/mL (Raji, U2932, RL) or 10 μg/ml (Ramos) rituximab and 20% human AB serum as a source of complement. Cell viability was assessed with PI staining. The survival of cells is presented as a percentage of control cells without antibody (mean± SEM). Statistical significance was determined using paired *t*-test, \*\**p<*0.01, \*\*\**p*<0.001, \*\*\*\**p*<0.0001 vs control WT cells. **(C)** Raji, U2932, Ramos, Daudi and LY7 cells were stably transduced with sgRNA (sg1 or sg3) silencing CD20 or with non-targeting control RNA (sgNTC). The levels of CD20 were assessed with flow cytometry. Representative overlays of CD20 MFI are presented. **(D)** Equal amounts of NTC and CD20 KO cells (sg1) were incubated for 1 h (Raji, Ramos, LY7, Daudi) or 4 h (U2932) with 100 μg/mL (Raji, U2932, LY7, Daudi) or 10 μg/ml (Ramos) rituximab and 20% human AB serum as a source of complement. Cell viability was assessed with PI staining. The survival of cells is presented as a percentage of control cells without antibody (mean± SEM). Statistical significance was determined using paired *t*-test, \*\*\**p*<0.001, \*\*\*\**p*<0.0001 vs controls. The experiments were repeated independently 4 times.

### CD20 KO cells are characterized with diminished CD37 levels

In order to investigate the consequences of CD20 downregulation, we characterized the surfaceome of three selected CD20 KO cell lines by multi-colour flow cytometry using the panel of mAbs against B-cell specific surface-restricted proteins (Fig. 2A). Although we observed changes in the levels of several antigens, the only universal change for all generated CD20 KO cell lines, except CD20 downregulation, was a decrease in the expression of CD37 (Fig. 2A). We further confirmed CD37 decrease in 6 consecutive CD20 KO cell lines using flow cytometry (Fig. 2B) and whole cell lysates using Western blotting (Fig. 2C, Suppl. Fig. 2A). Interestingly, we found no significant decrease in CD37 mRNA in CD20-defficient cells (Fig. 2D). Given the body of research showing synergistic action of anti-CD20 and anti-CD37 mAbs and suggesting close proximity of these two proteins (21,22), we sought to investigate if they form a complex in the cell membrane. Indeed, using Duolink Proximity Ligation flow cytometry-based assay which enables the detection of proteins within 40 nm distance (Fig. 2E), we observed the amplification generating green fluorescent signal in the presence of both anti-CD37 and anti-CD20 antibodies (Fig. 2F, Suppl. Fig. 2B). These results clearly demonstrated that CD20 and CD37 are close partners within the cell membrane. To understand their mutual regulation, we generated CD37 KO in Ramos cell line using the CRISPR-Cas9 approach, sorted out a pool and selected clones of CD37-negative cells (Suppl. Fig. 3A and B). To our surprise, we did not observe any reduction in CD20 levels associated with CD37 KO (Fig. 2G). These findings led us to conclude that CD20 and CD37 are closely associated on the cell membrane and that CD20 regulates the CD37 protein levels in a posttranscriptional mechanism.

**Figure 2.**
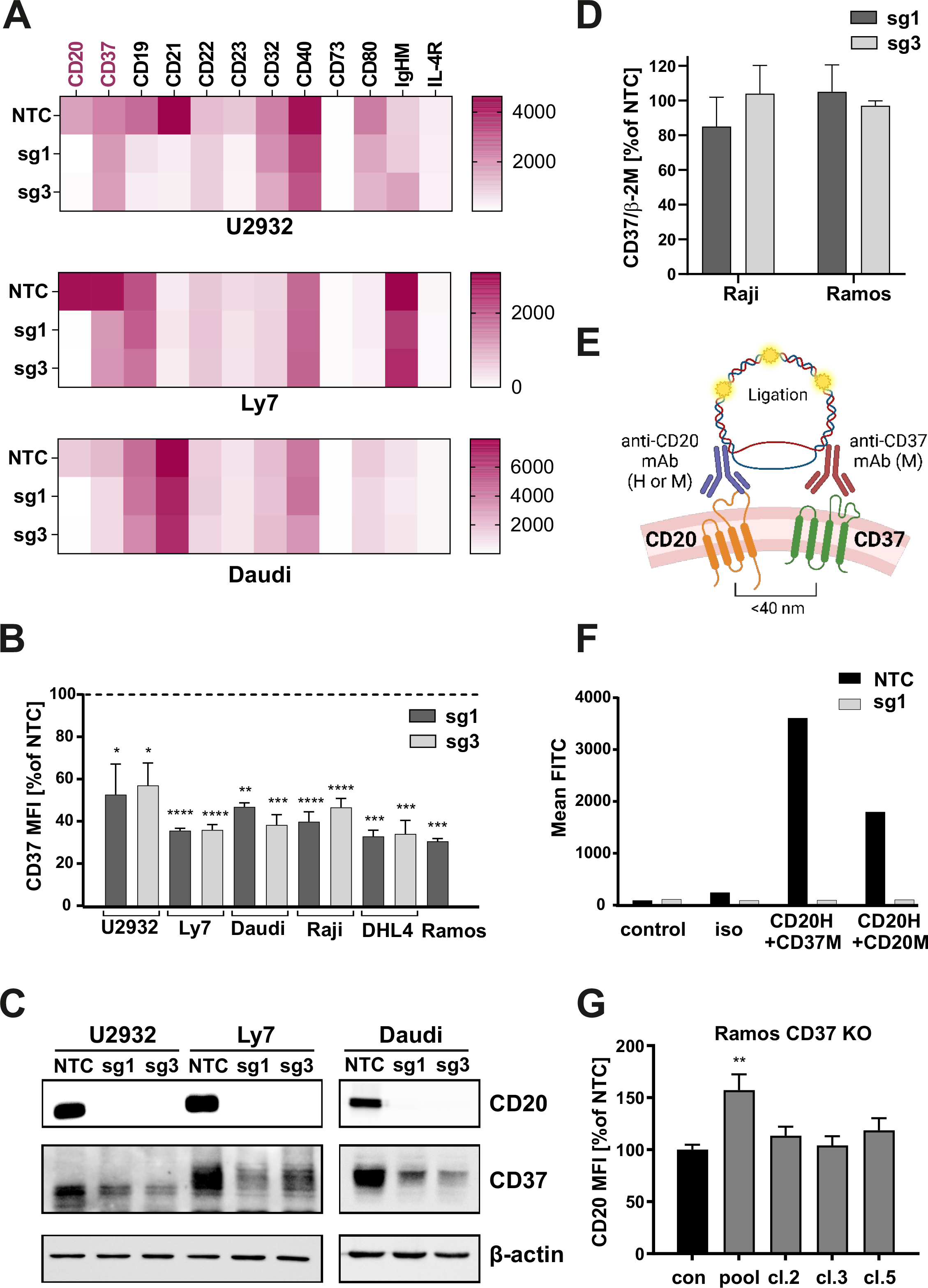
CD20 KO cells are characterized with diminished CD37 levels. **(A)** NTC and CD20 KO cells were stained with specific fluorochrome-conjugated antibodies targeting surface proteins typical for B-cells. The results are presented as a heatmaps. **(B)** NTC and CD20 KO cells were stained with FITC-conjugated anti-CD37 mAb. Dead cells were discriminated upon staining with PI. The results are presented as a percentage of MFI of NTC cells (mean ± SEM). Statistical significance was determined using 1-way ANOVA (NTC vs. sg1 or sg3) or with Welch’s *t*-test(NTC vs sg1) \**p*<0.05, \*\**p*<0.01, \*\*\**p*<0.001, \*\*\*\**p*<0.0001 vs. controls. **(C)** The levels of CD20 and CD37 were assessed with Western blotting in whole-cell lysates from NTC and CD20 KO cells. β-actin was used as a loading control. **(D)** cDNA from Raji and DHL4 NTC and CD20 KO – sg1 and sg3 cells was used for qRT-PCR amplification of CD37 and B2M. Relative reverse transcription quantitative polymerase chain reaction expression of CD37 gene was calculated by the user noninfluent second derivative method and shown as log-transformed target to reference ratio. B2M gene served as reference. **(E)** Proximity ligation experiment design. Cells of interest were incubated with saturating amounts of unlabelled human anti-CD20 mAb and mouse anti-CD37 mAb. Following incubation with anti-human and anti-mouse probes, in situ ligation and amplification, the formed protein complexes were analysed in flow cytometry. **(F)** PLA assay was performed in Raji NTC and sg1 cell lines to prove the formation of complex between CD20-CD37. The results are presented as MFI from a single experiment. The experiments were repeated independently 3 times. **(G)** Ramos WT cells were edited with CD37-targeting sgRNA, CD37-negative cells were sorted on a FACS sorter and single-cell clones were generated. Non-targeting sgRNA (NTC) was used as a negative control. CD20 expression was assessed by flow cytometry. The results are presented as a percentage of MFI of control cells (mean ± SEM). Statistical significance was determined using 1-way ANOVA, \*\**p*<0.01, \*\*\*\**p*<0.0001 vs controls.

### CD37 is downregulated in RR cells

To further explore the interaction of CD20 and CD37, this time in RR cells, we checked CD37 expression in RR cells, characterized by CD20 downregulation (Fig. 1A). In fact, CD37 was significantly down-regulated in all RR cell lines both on the cell surface (Fig. 3A) as well as in total cell lysates (Fig. 3B). Furthermore, in RR cells we have observed altered expression of several other antigens (Suppl. Fig. 4A) which reflects a multifaceted effect of RTX on the B-cell phenotype and existence of several mechanisms underlying the observed changes. Interestingly, the CD37 negative phenotype of RR cells persisted in vivo in mice inoculated with RR Ramos cells as demonstrated by the staining on the cells isolated from the spleen (Suppl. Fig. 4B). Moreover, in primary samples obtained from DLBCL patients before the start of first-line treatment, the expression of CD37 was heterogenous with approximately 50% of cases expressing CD37 at less than 5% of cells, as assessed by immunohistochemistry (Fig. 3C). This is in line with previously published observations (18). Furthermore, our results confirm the findings reported by Xu-Monette et al. showing the positive correlation between the presence of CD37 and PFS following R-CHOP therapy (Fig. 3D). Our clinicopathological correlations were limited to the newly diagnosed DLBCL cases (n=47) treated with R-CHOP in a first-line setting. Based on these experiments, we conclude that low/decreased CD37 expression can be relatively frequent in patients. Since CD37 is explored as a therapeutic target, in the subsequent steps, we investigated the consequences of decreased CD37 levels on the efficacy of various forms of therapy targeting CD37.

**Figure 3.**
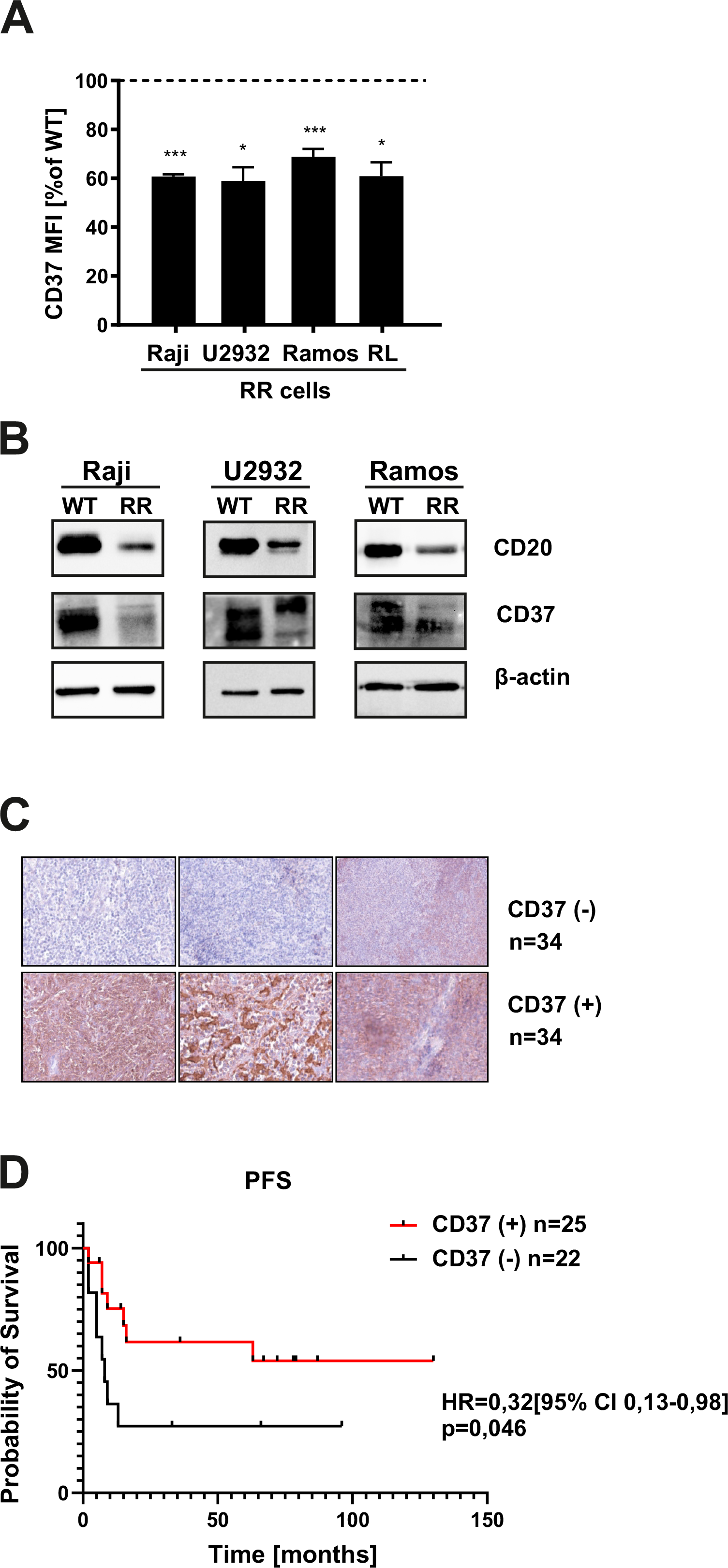
CD37 is downregulated in RR cells. **(A)** WT and RR cells were stained with FITC-conjugated anti-CD37 mAb. The viability was assessed using PI. The results are presented as a percentage of MFI of control cells (mean ± SEM). Statistical significance was determined with Welch’s *t*-test, \**p*<0.05, \*\*\**p*<0.001. The experiments were repeated independently 4 times. **(B)** The levels of CD20 and CD37 were assessed with Western blotting in whole-cell lysates. β-actin was used as loading control. The experiments were repeated independently 3 times. **(C)** CD37 expression was evaluated in 68 primary paraffin-embedded samples by a pathologist blinded to the clinical data of the patients. Representative stainings for CD37-positive and CD37-negative stainings are shown. **(D)** Kaplan-Meier curves were plotted for PFS in CD37- and CD37+ patients. The survival curves were compared with Peto and Peto’s generalized Wilcoxon test.

### CD37 decrease leads to impaired efficacy of CD37 mAbs-mediated CDC, which can be restored using lysosome inhibitors

A decrease in the target antigen directly translates into impaired efficacy of mAbs that exert cytotoxicity in complement-dependent mechanism (23,24). Therefore, we have asked the question if decreased CD37 levels in RR and CD20 KO cells influence the efficacy of a CDC-inducing anti-CD37 mAb. We observed impaired efficacy of anti-CD37 mAb in CD20 KO and RR cells (Fig. 4A and B, respectively). This effect was clearly visible despite utilizing a mAb with the E430G mutation, which promotes more efficient IgG hexamer formation through intermolecular Fc-Fc interactions after cell surface antigen binding (25,26). While in CD20 KO cells a decreased efficacy of anti-CD37 mAb may be solely attributed to decreased CD37 expression, in RR cells it may be the result of both decreased CD37 expression and an increase in complement inhibitor protein CD59 (Suppl. Fig. 4C).

**Figure 4.**
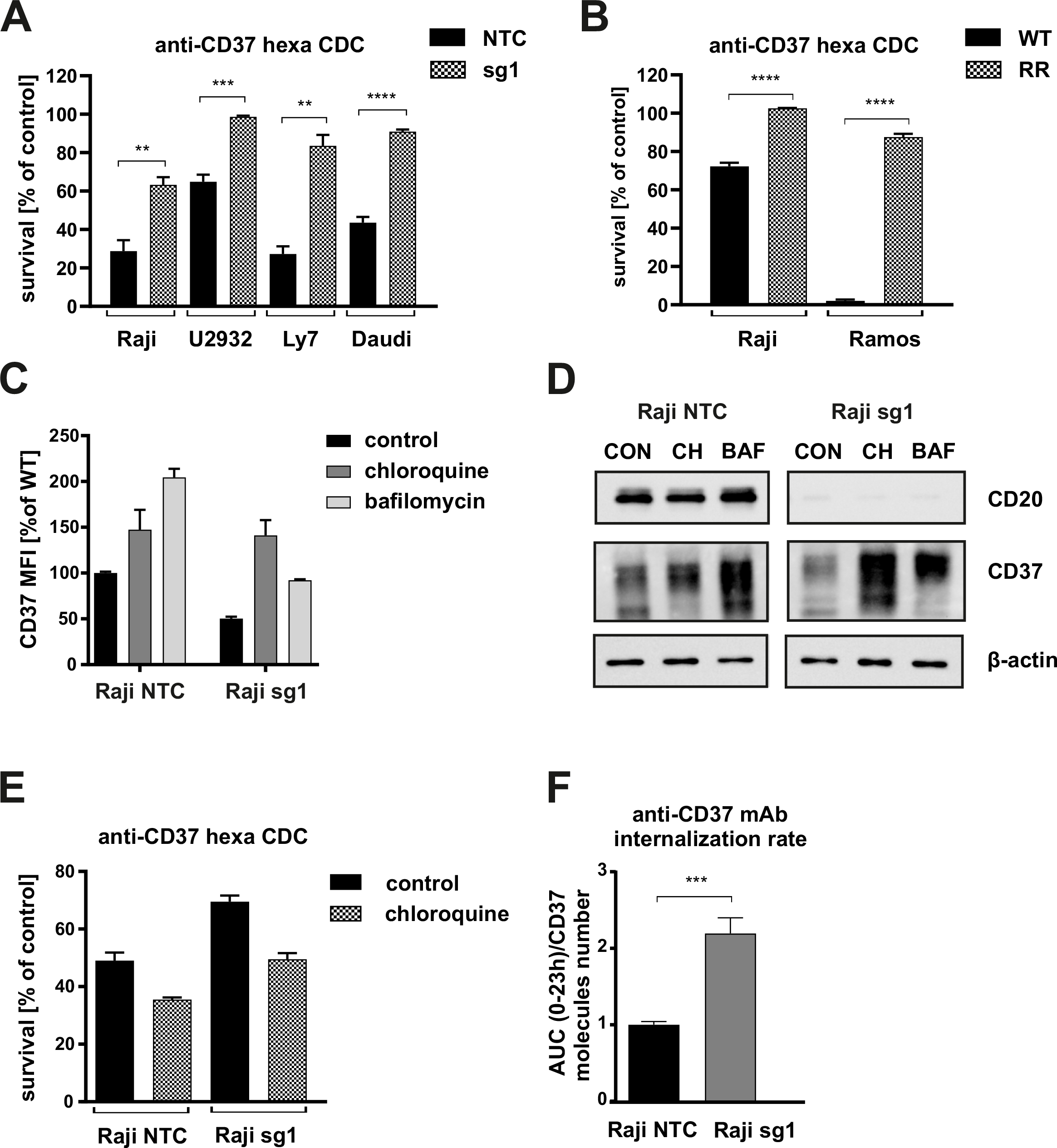
CD37 decrease leads to impaired efficacy of anti-CD37 mAbs-mediated CDC, which can be restored using lysosome inhibitors. Equal amounts of NTC and CD20 KO cells **(A)** and WT and RR Raji cells **(B)** were incubated for 1 h (Raji, Ramos, Daudi, LY7) or 4 h (U2932) with 10 μg/ml anti-CD37 mAb and 20% human AB serum as a source of complement. Cell viability was assessed with PI staining. The survival of cells is presented as a percentage of control cells without antibody (mean± SEM). Statistical significance was determined using paired *t*-test, \*\**p*<0.01, \*\*\**p*<0.001, \*\*\*\**p*<0 .0001 vs controls. **(C)** Equal amounts of NTC and CD20 KO cells were incubated for 24 h with non-toxic concentrations of lysosome inhibiting agents – 100 μM chloroquine and 20 nM bafilomycin. Thereafter, the cells were stained with FITC-conjugated anti-CD37 mAb. The results are presented as a percentage of MFI of control cells (mean± SD). Representative results are shown. **(D)** The levels of CD20 and CD37 were assessed with Western blotting in whole-cell lysates. β-actin was used as loading control. **(E)** Equal amounts of NTC and CD20 KO Raji cells preincubated for 36 h with 100 μM chloroquine were treated for 1 h with anti-CD37 mAb (5-10 μg/ml) and 20% human AB serum as a source of complement. Cell viability was assessed with PI staining. The survival of cells is presented as a percentage of control cells without antibody (mean± SEM). *(F)* Internalization rate of mouse anti-human anti-CD37 mAb was determined in NTC and sg1 cells incubated in IncuCyte with equal amounts of mAb conjugated to pH sensitive dye activated in lysosome. The internalization rate is shown as fluorescence AUC normalized to number of CD37 molecules per cell.

Since CD20 and CD37 are interrelated in the plasma membrane, we hypothesized that the mechanisms of their interaction are dependent on the internalization, trafficking and degradation processes. More specifically, we assumed that CD37 endocytosis is accelerated in the cells lacking CD20 and that the presence of CD20 stabilizes CD37 within the cell membrane. To test this assumption, we have preincubated NTC and CD20 KO cells with non-toxic concentrations of compounds known to block lysosome-dependent degradation i.e. chloroquine and bafilomycin A. The used inhibitors up-regulated CD37 levels in both NTC as well as in CD20 KO cells, where they increased CD37 levels up to the values observed in NTC cells (Fig. 4C). Increase in CD20 and CD37 levels can also be observed in whole-cell lysates of these cells (Fig. 4D). Accordingly, inhibition of lysosomal degradation increased the efficacy of anti-CD37 mAbs against CD20 KO cells in CDC cytotoxicity assay (Fig. 4E). All in all, our results show that CD37 can be degraded by lysosome and the blockage of lysosome can be used as an approach to increase the sensitivity of tumor cells with CD20 KO to anti-CD37 mAbs.

### CD20 KO cells show an increased internalization rate of anti-CD37 mAb

In the further steps, we investigated the potential of the use of anti-CD37 antibody-drug conjugates (ADCs) in the cells with decreased CD20 levels. This treatment approach involves the use of surface antigen-targeted mAbs to deliver cytotoxic payloads into tumor cells upon conjugate internalization through endocytosis. In our experiments, we investigated the internalization rate of anti-CD37 mAb in NTC and CD20 KO cells. We were able to observe increased internalization rate of a mouse anti-human anti-CD37 antibody. In this experiment, Raji NTC and CD20 KO cells were incubated with equal amounts of mouse anti-human anti-CD37 mAb conjugated with pH-sensitive dye, which becomes activated in lysosomes, for 23h in IncuCyte, and the images were registered every 20 min. Fluorescence AUC was normalized to the cell area (Suppl Fig. 4A). As CD20 KO cells were characterized with decreased CD37 levels, Quantibrite measurement was performed (Suppl. Fig. 4C) and the AUC normalized to the number of CD37 molecules per cell (Fig. 4F). These results suggest an increased internalization in CD20 KO cells as one of the potential mechanisms involved in the downregulation of CD37 that could be exploited therapeutically with anti-CD37 ADCs.

### CD37 decrease does not hamper CAR T cells’ cytotoxicity

Since CD37 has recently been designated as a suitable target for CAR-directed therapy (11,12), we then asked the question if a substantial decrease in its levels would hamper the efficacy of CD37 CARs as well. To address this, we performed CD37 CAR-T cytotoxic assays against both RR and CD20 KO cell lines as targets in two effector-to-target (E:T) ratios. We have discovered that despite significantly diminished CD37 levels in 6 different cell lines (either RRCLs or CD20 KO cells), the cytotoxicity of the tested CAR T cells was not significantly reduced (Fig. 5A and B). At this stage, we concluded that CD37, although diminished upon downregulation of CD20, remains an attractive target for cell-based therapies and could be further explored even in RTX-pretreated patients.

**Figure 5.**
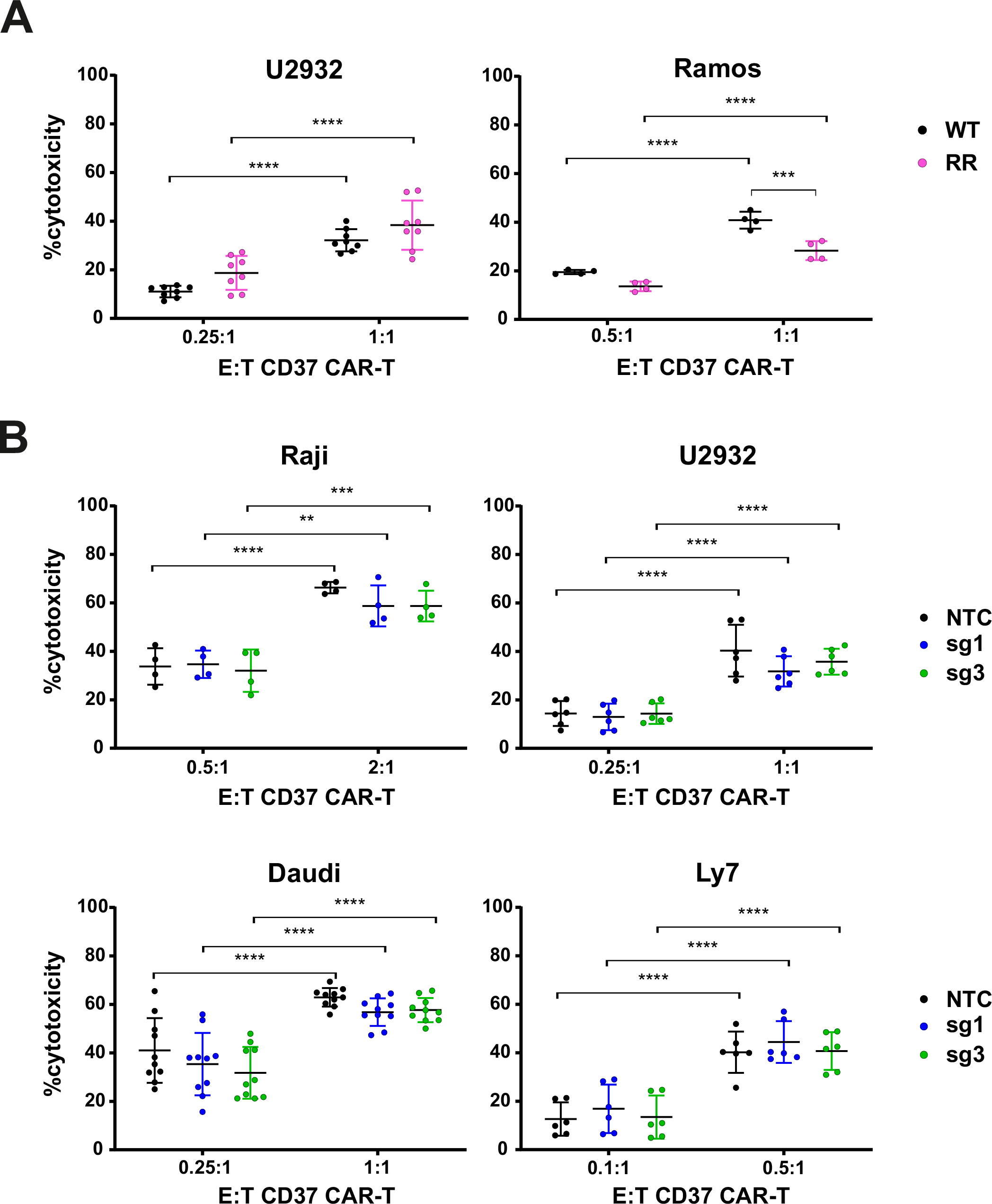
CD37 decrease does not hamper CAR T cells’ cytotoxicity. Equal amounts of **(A)** WT and rituximab-resistant (RR) or **(B)** NTC and sg1 cells were stained with CellTrace™ Violet and incubated for 16 h with various effector to target (E:T) ratios of CD37 CAR-T cells. Cell viability was assessed with flow cytometry following incubation PI staining. Data represent mean ± SD of % cytotoxicity of CD37 CAR-T cells after subtracting cytotoxicity of unmodified T-cells. Statistical significance was determined using 2-way ANOVA with Tukey’s post hoc test. Experiments were repeated at least 4 times.

## Discussion

Novel treatment modalities for r/r DLBCL patients are intensively pursued. However, little is known about the mechanisms of primary resistance and molecular changes induced by the treatment on the phenotype of lymphoma cells despite the clear unfavourable prognosis of patients progressing during or shortly after first-line immunochemotherapy (27). In the recent few years, the therapeutic armamentarium for aggressive lymphomas has expanded thanks to registrations of second-line treatments that finally offer an alternative to auto-HSCT, namely anti-CD19 CAR T cells, novel mAbs-based regimens: tafasitamab plus lenalidomide and polatuzumab vedotin and novel promising agents such as novel ADCs and bispecific mAbs tested in clinical trials (28).

CD37, a tetraspanin predominantly expressed on B cells is currently being explored as a target for mAbs, ADCs and CAR T cells. The results of our work show that strategies targeting CD37 need to be carefully implemented in r/r B-cell lymphoma patients. Using CD20 KO cells, we identified the association between CD37 and CD20, with the expression of the latter playing a crucial protective role in preventing CD37 from undergoing endocytosis as suggested by our results in CD20 KO cells where lysosome inhibition increased reduced CD37 levels to the levels seen in control (NTC) cells. This discovery has several implications. First of all, we demonstrate that decreased CD37 expression correlates with reduced efficacy of anti-CD37 mAbs in a complement-dependent mechanism, where the expression of the target antigen determines the cytotoxic effect. However, this limitation may be encompassed by the use of bi-specific antibodies e.g. targeting CD20 and CD37 that show superior efficacy to anti-CD37 mAbs alone (21) or bi-paratopic mAbs (24,29).

On the other hand, we hypothesize that CD20 loss facilitates CD37 endocytosis and it may translate to increased efficacy of anti-CD37 ADCs that have recently been presented as a highly efficient strategy in B-cell malignancies in vitro and in murine models (30). The efficacy of ADCs relies on the endocytosis of the target antigen-ADC complex and the release of the cytotoxic payload to the cytoplasm. It has already been reported that rituximab synergizes with naratuximab emtansine by increasing the endocytosis of CD37 (22). Our results suggest that anti-CD37 ADCs may be effective even in the case of CD20-negative cells in r/r RTX-treated patients. Finally, we show that decreased CD37 levels (up to 50% in our models) do not influence the efficacy of CD37-targeted CARs and delineate CD37 CARs as an effective option for CD20-negative RTX-pretreated patients. However, approximately 50% of DLBCL patients have undetectable CD37 levels in immunohistochemistry (IHC), the reason behind this phenomenon being attributed at least partially to low levels of IRF8, a newly identified transcriptional regulator of CD37 expression(31).

Moreover, our study sheds new light on the regulation of CD37 expression. Decrease in CD37 expression has been reported to affect patients’ prognosis negatively, and according to the studies by other groups (31) and ours, up to 50% of DLBCL patients show undetectable CD37 protein expression by immunohistochemistry. However, it needs to be emphasized that in the IHC stainings performed by us and others a clone of anti-CD37 mAb (2B8) used does not correspond to the clone used for CAR construct (HH1 clone). As mentioned before r/r patients are not routinely rebiopsed therefore not much is known about the changes in surfaceome. Also, IHC may be an imperfect tool to study such changes as demonstrated by the recent study (32), where decreases in CD20 levels were observed in flow cytometry in the samples where no decrease in CD20 by IHC has been noted. By now, CD37 loss has been attributed to decreased expression a transcriptional factor IRF8 (31). However, as demonstrated by the study by Elfrink et al., up to 40% of IRF8 low DLBCL samples still express detectable CD37 in IHC (31) suggesting that there are additional factors regulating CD37 expression. Tetraspanins play an important role in the organization of the cellular membrane (33), forming clusters that overlap each other. The CD37 cluster has been shown to overlap with CD81 and CD82 cluster (33), while the latter has been reported to form a supramolecular complex with MHC class I, MHC class II, CD53, and CD20 (34). Our results suggest that a decrease in CD20 levels, a protein not belonging to the tetraspanin family but to MS4A family that shows some characteristics in the structure, may induce changes in the composition of the cellular membrane and promote the degradation of CD37. Other members of MS4A family have already been demonstrated to govern the endocytosis process i.e. MS4A4A by directing the trafficking of KIT regulates its signaling(35) and MS4A3 promotes endocytosis of common β-chain (βc) cytokine receptors (36).

All in all, our study adds new knowledge on the current state of the art on the regulation of CD37, which in the light of the recent findings (11,12,18,24,29,30) constitutes an attractive therapeutic target for immunotherapies. The results of our study once again prove that CD37 is highly variable in DLBCL patients. Finally, we show that immunotherapy using anti-CD37 CAR T cells can be an effective option despite low levels of CD37 protein.

## Acknowledgments

The authors would like to thank Prof. F.Hernandez-Iizaliturri, Prof. M. Czuczman and Cory Mavis from Roswell Park, NY, US for providing RR cell lines and Genmab, Utrecht, the Netherlands for providing anti-CD37 mAb.

## Authors’ contribution

M.B. designed the study, performed and analyzed experiments and wrote the manuscript, A.K. performed and analyzed experiments and was involved in manuscript preparation; M.K., A.S., J.D., M.Ku., M.S., L.D., K.M., K.F., C.F., M.P., I.B., A.G-J., C.L.G. performed and analyzed experiments, J.B. provided primary material and analyzed patients’ data, A.S-C. performed immunohistochemistry analysis, E.M.I. and S.W. provided CAR CD37 construct and were involved in manuscript preparation, M.P., M.F. and M.P-S. provided expert advice and guidance throughout the study, M.W. designed the study, wrote the manuscript, prepared the figures and provided expert guidance throughout the study; all authors reviewed the manuscript.

## Data Availability Statement

Raw data will be made available upon reasonable request to the corresponding author.

