## Supplement_material for "CD20 expression regulates CD37 levels in B-cell lymphoma – implications for immunotherapies"

### Supplemental Figure 1

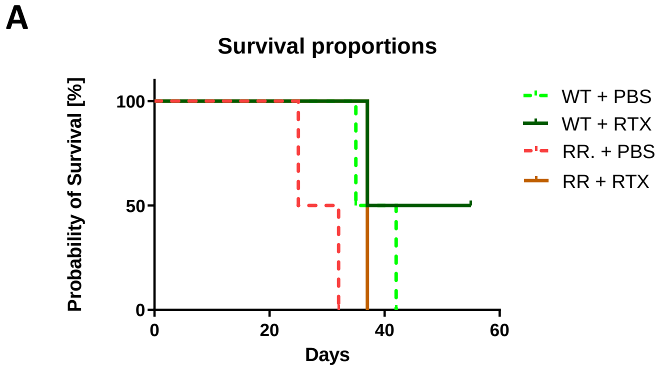


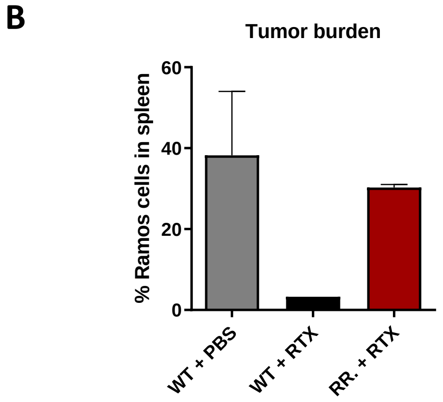

### Supplemental Figure 2

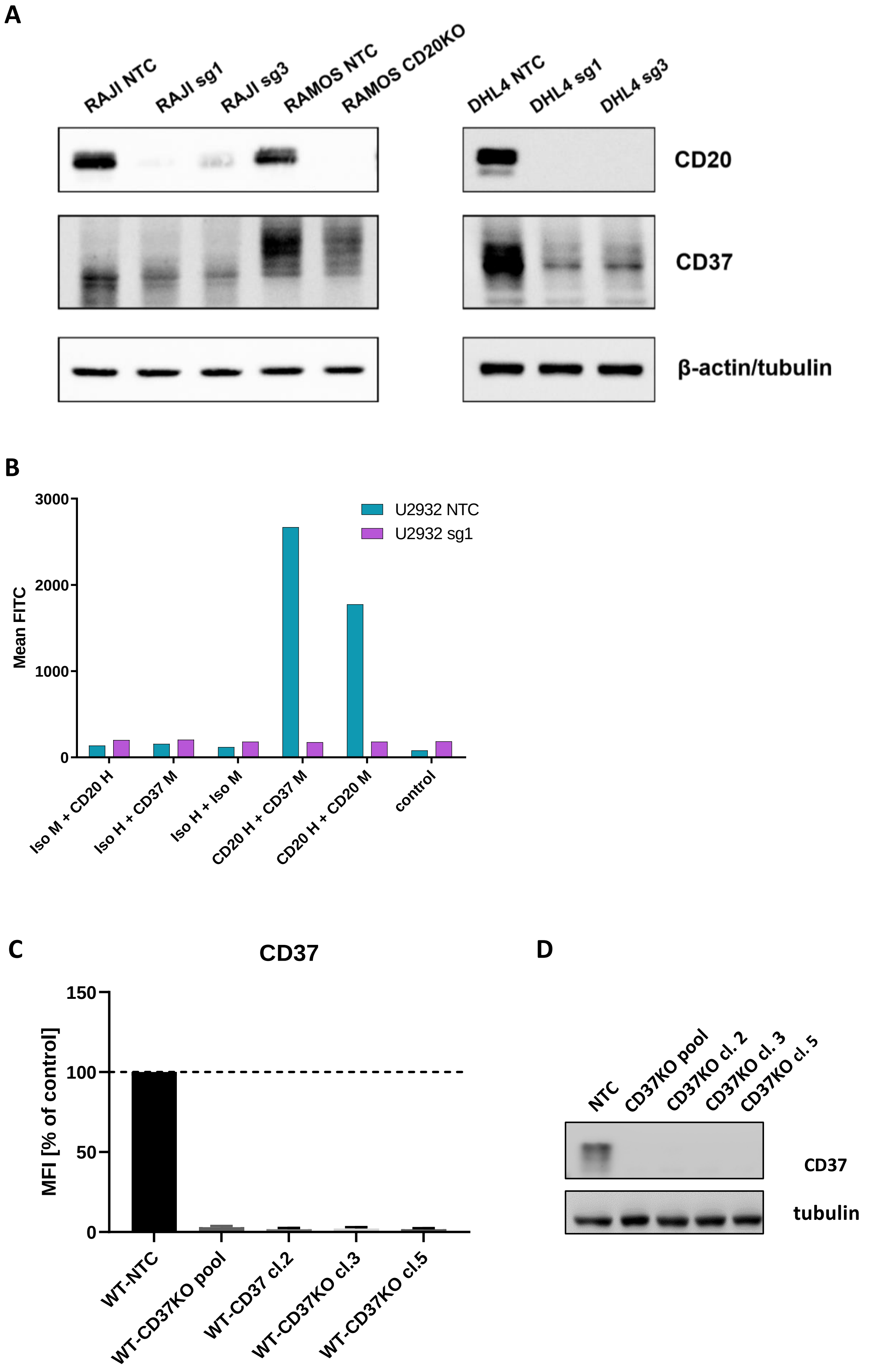

### Supplemental Figure 3

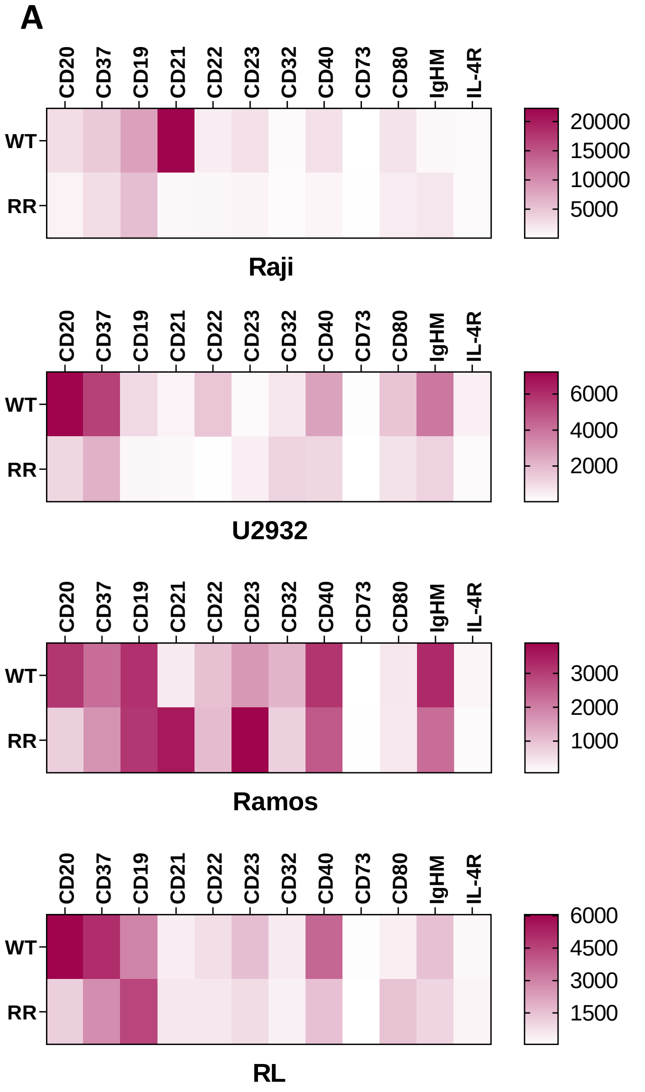


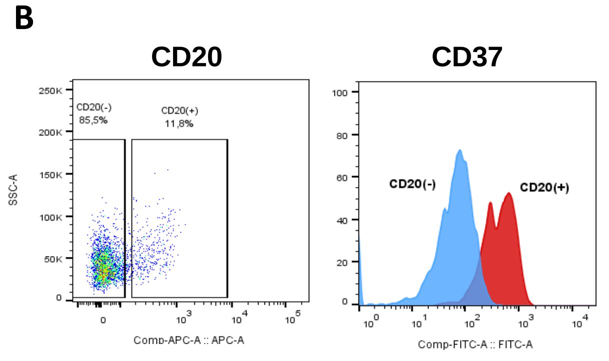


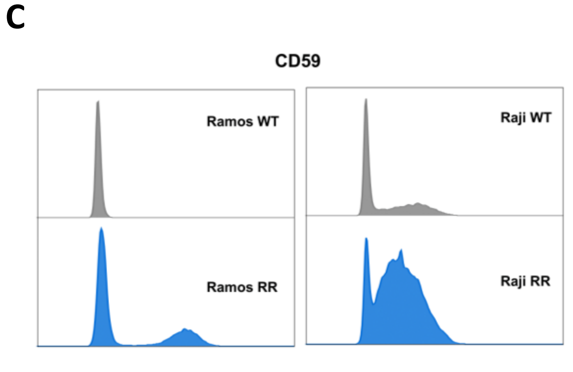

### Supplemental Figure 4

A

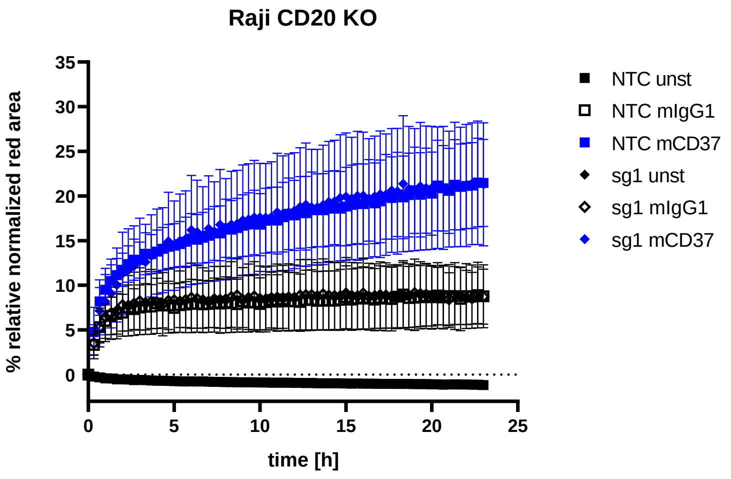


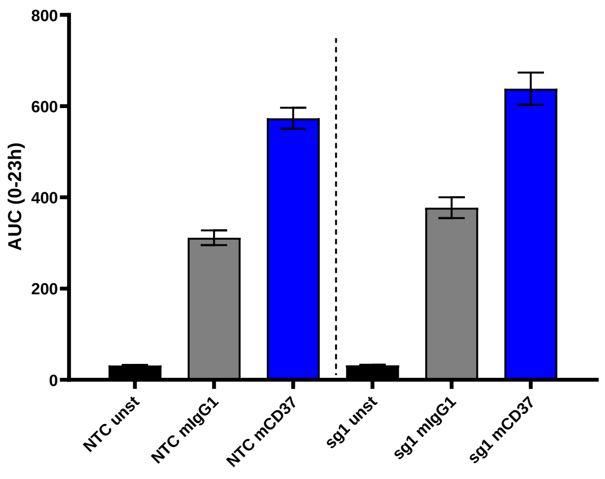


C

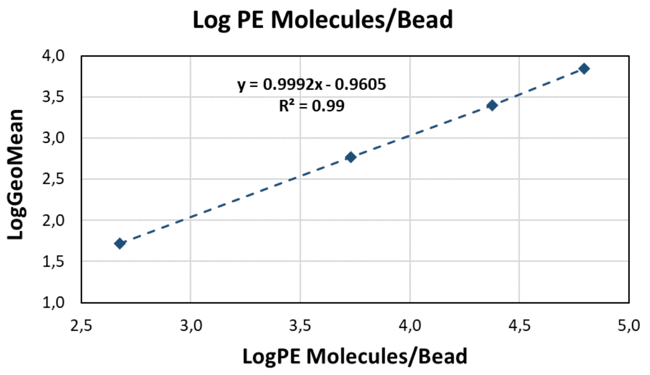


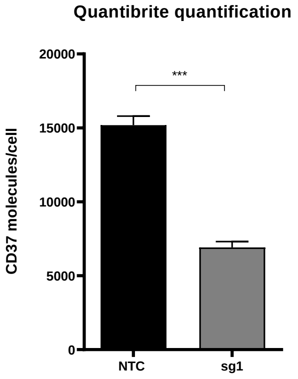

**Supplemental Table 1**

| **Antigen** | **Clone** |
|---|---|
| **CD20** | **L27** |
| **CD21** | **HB5** |
| **CD19** | **HIB19** |
| **CD40** | **5C3** |
| **CD37** | **M-B271** |
| **CD32** | **6C4** |
| **CD80** | **L307.4** |
| **CD22** | **4KB128** |
| **IgHM** | **SA-DA4** |
| **CD23** | **EBVCS2** |
| **CD74** | **5-329** |
| **IL-4R** | **25463** |
| **CD73** | **AD2** |

**Supplemental Table 2**

| | **sequence** |
|---|---|
| **CD20KO_sg1** | GGATCATCAGAAGACCCCCC |
| **CD20KO_sg3** | CAGCAACGGAGAAAAACTCC |
| **CD37KO** | TTTGCCACACAGATCACCCT |
| **NTC** | CACCGGGGCGAGGAGCTGTTCACCG |

Supplemental Figure 1. RR cells' growth *in vivo*

2 million Ramos cells (WT or RTX-resistant) were injected in 100ul PBS into the tail vein of NOD-Scid-IL2Rgamma (NSG) immunodeficient mice. Four weeks later, when the mice started to exhibit first signs of disease, the mice were randomly divided into two groups and treated either with RTX (100 mg/kg)

or with PBS, every 3 days intraperitoneally. At the signs of mouse distress due to progressive disease, the animals were sacrificed, their spleen was removed, mashed up and splenocytes were collected. **(A)** Survival curves. **(B)** Tumor burden was assessed by analyzing the percentage of CD45 positive cells in the spleen.

Supplemental Figure 2. CD37 is downregulated in CD20KO cells

(A) The levels of CD20 and CD37 were assessed with Western blotting in whole-cell lysates from NTC and CD20 KO cells. β-actin or tubulin were used as loading controls. (B) PLA assay was performed in U2923 NTC and sg1 cell lines to prove the formation of complex between CD20-CD37. The results are presented as MFI from a single experiment. (C-D) Ramos WT cells were modified with NTC and CD37 KO encoding constructs. CD37 KO was confirmed with flow cytometry and Western blotting.

Supplemental Figure 3. RR cells are characterized with CD37 downregulation

(A) WT and RR cells were stained with specific fluorochrome-conjugated antibodies targeting surface proteins typical for B-cells. The results are presented as a heatmap. (B) Spleens from mice injected with Ramos RR cells were isolated. CD20 and CD37 expression were analyzed in flow cytometry. (C) CD59 expression was assessed in WT and RR Ramos and Raji cells in flow cytometry.

Supplemental Figure 4. Anti-CD37 mAb is efficiently internalized in both NTC and CD20KO cells

Raji NTC and sg1 cells were incubated with mouse anti-human anti-CD37 mAb and isotype control mAb conjugated with pH-sensitive fluorescence dye in the IncuCyte high-throughput fluorescence microscope system. **(A)** The fluorescence images of each well were created every 20 minutes for 23 h. Fluorescence curves of each well are demonstrated. Internalization was defined as relative normalized red area in percent (%) over the time: red fluorescence area (Total Red Area per Image) was normalized (divided by) to the total cell area (phase confluence per Image) in percent (%) relative to time point 0 h. **(B)** Area under the curve (AUC) of fluorescence curves 0-23 h. **(C)** Quantibrite quantification of CD37 expression was performed in NTC and sg1 Raji cells. The results are presented as CD37 molecules/cell ± SEM. The experiments were repeated 3 times.

#### Supplemental Methods

##### Reagents

Rituximab and obinutuzumab were purchased from Roche and anti-CD37 E430G antibody was a generous gift from Genmab. Ibritumomab and trastuzumab that were used in PLA assays were purchased from ProteoGenix. Chloroquine, bafilomycin A and propidium iodide (PI) were purchased from Sigma Aldrich. Human serum was purchased from Blood Collection Center in Warsaw.

##### Staining of surface antigens

Cells were stained as described earlier in [19]. Cells were incubated with saturating amounts of fluorochrome-conjugated Abs or IgG1 isotype controls (Suppl. Table 1) for 30 min in RT. For viability staining the cells were either incubated with Fixable Viability Stain 510 (BD Horizon) prior to antigen staining or they were resuspended in PBS supplemented with 4 µg/ml propidium iodide (PI) before analysis.

##### Plasmids and transductions

Sequences for CD20 KO -sg1, sg3, CD37KO as well as sgEGFP (further referred to as NTC) were generated with oligonucleotide pairs selected from Brunello CRISP Knockout Pooled Library (Supplementary Table 2) and cloned into pLenti-CRISPRv2, a gift from Feng Zhang, Addgene plasmid #52961. HEK 293T cells seeded into 10 cm plates were used for the production of a replication-incompetent lentivirus. CAR CD37 plasmid was generated as described in [12]. All the plasmids were sequenced using standard Sanger sequencing by oligo.pl (Institute of Biochemistry and Biophysics, Warsaw, Poland). Lentiviral and retroviral particles were produced as described previously in [19]. Cells containing gene-of-interest were selected using puromycin and sorted for CD20-negative populations using BD FACS Aria.

CD37KO sequence was generated as described above (Supplementary Table 2) and cloned into LentiGuide-puro vector (a gift from Feng Zhang, Addgene plasmid # 52963), followed by lentiviral infection of target cells and their puromycin selection. Target cells were infected with lentiCas9-blast (a gift from Feng Zhang, Addgene plasmid # 52962) and selected with blasticidin. The CD37-deleted cells were then sorted out by FACS sorting and expanded. To isolate single-cell clones, CD37-negative cells were sorted out into 96-well plate as 1 cell per well and allowed to expand.

In vivo experiments

2 million Ramos cells (WT or RR) were injected in 100 ul PBS into the tail vein of NOD-Scid-IL2Rgamma (NSG) immunodeficient mice. Four weeks later, when the mice started to exhibit first signs of disease, the mice were randomly divided into two groups and treated either with RTX (100 mg/kg) or with PBS, every 3 days intraperitoneally. At the signs of mouse distress due to progressive disease, the animals were sacrificed, their spleen was removed, mashed up and splenocytes were collected. Cells were stained with CD20-APC and CD37-FITC antibodies and analyzed by flow cytometry.

**Cytotoxicity assays**

CDC assay was performed as described earlier [19]. Briefly, lymphoma cells were incubated for 1 or 4 h (depending on the cell line's sensitivity) with RTX or anti-CD37 E430G antibody in the presence of 20% human serum. For CAR-T cytotoxicity assays, target tumor cells were stained with 0,5 uM Cell Trace Violet (Thermo Scientific) and seeded with CAR-T cells and unmodified T-cells at different effector to target (E:T) ratios for 16 h. The viability of tumor cells was analysed with flow cytometry following PI staining.
